# Aberrant and Ectopic Cell Populations of the Fibrotic Pushing Border in Restrictive Allograft Syndrome after Lung Transplantation

**DOI:** 10.1101/2024.06.04.597358

**Authors:** Lena M. Leiber, Leonard Christian, Lavinia Neubert, Jannik Ruwisch, Hande Yilmaz, Edith K. J. Plucinski, Linus Langer, Jan C. Kamp, Mark Greer, Bernd Haermeyer, Mark Kühnel, Christopher Werlein, Aurelien Justet, Anke K. Bergmann, Matthias Ballmaier, Jawad Salman, Lars Knudsen, Ulrich Martin, Bart Vanaudenaerde, Ali Önder Yildirim, Fabio Ius, Florian Laenger, Tobias Welte, Christine Falk, Naftali Kaminski, Danny D. Jonigk, Jens Gottlieb, Jonas C. Schupp

**Affiliations:** Department of Respiratory Medicine and Infectious Diseases, Hannover Medical School, Hannover, Germany; Biomedical Research in Endstage and Obstructive Lung Disease Hannover, German Center for Lung Research, Hannover, Germany; Institute of Pathology, Hannover Medical School, Hannover, Germany; Institute of Human Genetics, Hannover Medical School, Hannover, Germany; Institute of Pathology, RWTH Aachen, Aachen, Germany; Service de Pneumologie, CHU de Caen, Centre de compétence des maladies pulmonaires rares, ISTCT, UMR6030-CNRS-CEA-Université de Caen, Caen, France; Central Research Facility Cell Sorting, Hannover Medical School, Hannover, Germany; Clinic for cardiac, thoracic, transplant and vascular surgery, Hannover Medical School, Hannover, Germany; Institute for Functional and Applied Anatomy, Hannover Medical School, Hannover, Germany; Leibniz Research Laboratories for Biotechnology and Artificial Organs, Lower Saxony Center for Biomedical Engineering, Implant Research and Development /Department of Cardiothoracic, Transplantation and Vascular Surgery, Hannover Medical School, Carl-Neuberg-Str. 1, 30625 Hannover, Germany; Department of Chronic Diseases and Metabolism, KU Leuven, Leuven, Belgium; Institute of Lung Health and Immunity (LHI), Helmholtz Zentrum Muenchen, Institute of Lung Biology and Disease; Munich, Germany; Institute of Transplant Immunology, Hannover Medical School, Hannover, Germany; Section of Pulmonary, Critical Care and Sleep Medicine, Yale School of Medicine, New Haven, CT, United States; Department of Clinical Airway Research, Fraunhofer Institute for Toxicology and Experimental Medicine (ITEM), Hannover, Germany

## Abstract

**Rationale:** Restrictive allograft syndrome (RAS) is a major cause of mortality in patients following lung transplantation due to rapid progressive fibrosis in the pulmonary graft. We have only limited knowledge of the cellular and molecular mechanisms that characterize the fibrosis in the RAS lung.

**Objective:** To elucidate cellularly-resolved transcriptomic and histologic characteristics of the structural cells in human RAS lungs.

**Methods:** Single-nuclei RNA-sequencing was performed in peripheral lung tissues from 15 RAS patients undergoing lung re-transplantation, and from 9 healthy control lungs. Findings were validated and complemented by various histologic techniques, including immunofluorescence, RNAscope, combined Elastica van Gieson-immunohistochemistry stains, and micro-CT scans.

**Measurement and Main results:** Differential gene expression analysis of our single-nuclei RNA-sequencing data revealed in human RAS lungs previously undescribed and uniquely distributed aberrant basaloid cells, ectopic *COL15A1+* vascular endothelial cells, and *CTHRC1+* fibrotic fibroblasts, all first characterized in idiopathic pulmonary fibrosis (IPF). In contrast to IPF, RAS lacks the cellular equivalent of bronchiolization. Histologic stains confirmed our transcriptomic discoveries and disclosed distinctive distribution patterns: Aberrant basaloid cells are primarily localized at the edge of the fibrotic pushing border, forming together with the juxtaposed *CTHRC1+* fibrotic fibroblasts the fibrotic niche of alveolar fibroelastosis (AFE), the histopathological hallmark in RAS lungs. On the endothelial side, *PRX+* alveolar microvasculature is lost in AFE areas. Micro-CT scans revealed that blood supply, now facilitated by expanded and ectopic *COL15A1+* VE cells, changes from pulmonary to systemic perfusion. Last, our data reveals potential therapeutically-modifiable expression patterns in RAS, including genes coding for the integrin subunits αvβ6, activators of TGFβ.

**Conclusion:** Considering the marked clinical, histologic and etiologic dissimilarities of RAS and IPF, our snRNAseq study revealed a surprising general principle of cellular and molecular pathogenesis in the fibrosing lung: the entity-spanning composition of the fibrotic niche by a) aberrant basaloid cells localized at the fibrotic pushing border, b) ectopic *COL15A1+* vascular ECs and c) effector *CTHRC1+* fibrotic fibroblasts. This general principle justifies a flexible but cellular pathogenesis-guided transferability of potential therapeutic approaches between progressive fibrotic lung diseases.

## INTRODUCTION

Chronic lung allograft dysfunction (CLAD) after lung transplantation (LuTx) is an aggressive remodelling process, related to a multitude of causes, including allo- and auto-immunity, infection, aspiration and ischemia, and manifesting as restrictive and/or obstructive phenotypes, depending on the pulmonary compartments involved and the predominant injury pattern.(1,2) Due to lacking therapy options, CLAD remains a major cause of post-LuTx morbidity, necessity of re-transplantation (ReTx), and mortality after LuTx. Up to 30% of CLAD patients are diagnosed with a restrictive phenotype, called restrictive allograft syndrome (RAS).(2,3) RAS is associated with a worse prognosis compared to the obstructive phenotype, bronchiolitis obliterans syndrome (BOS) (survival rate after diagnosis: 6 - 18 months (RAS) vs 3 - 5 years (BOS)).(2,4–7) The pathogenesis of RAS remains enigmatic while some potential risk factors such as acute cellular rejection (ACR), donor-specific antibodies (DSA) and late-onset diffuse alveolar damage (DAD) have been identified. Large parts of today‘s vastly incomplete knowledge concerning the pathogenesis and pathognomonic changes of RAS and CLAD originates from bronchoalveolar lavage fluid (BALF) data and blood analyses or animal models(5,7–9).

RAS lung tissue is characterized histologically by the juxtaposition of primarily sub-pleural and perivenular dense fibrotic areas, termed alveolar fibroelastosis (AFE), which sharply transition to alveoli with an inconspicuous architecture.(5,10,11) AFE itself is a unique histological pattern characterized by the collagenous obliteration of alveoli, and de-epitheliazation and elastosis of alveolar walls with only scant inflammation(10). In this manuscript, we will strictly refer to the visual, sharp border between areas of dense fibrosis/AFE and structurally intact alveolar lung tissue as ‘fibrotic pushing border’. AFE progresses centripetally, predominantly starting from thickened pleura in the apex of the upper lobes.

Within this study, we performed the first cellularly-resolved transcriptomic analysis of human RAS lungs to elucidate its transcriptomic characteristics in relation to the histopathological characteristics of the pulmonary injury pattern(s) present. Single-nuclei RNA-sequencing (snRNA-seq) data enabled us to identify aberrant epithelial, ectopic endothelial and fibrotic fibroblast populations which were previously undiscovered in RAS. Immunhistofluorescent (IHF), RNAscope, and combined Elastica van Gieson-immunohistochemistry (EvG-IHC) stains, and micro-CT scans provide insights in the pathogenetic characteristics of RAS, revealed specific distribution patterns of disease-enriched cell populations and enabled a complementary 3D-structural analysis of our snRNA-seq-based findings (Fig. 1A).

**Figure 1.**
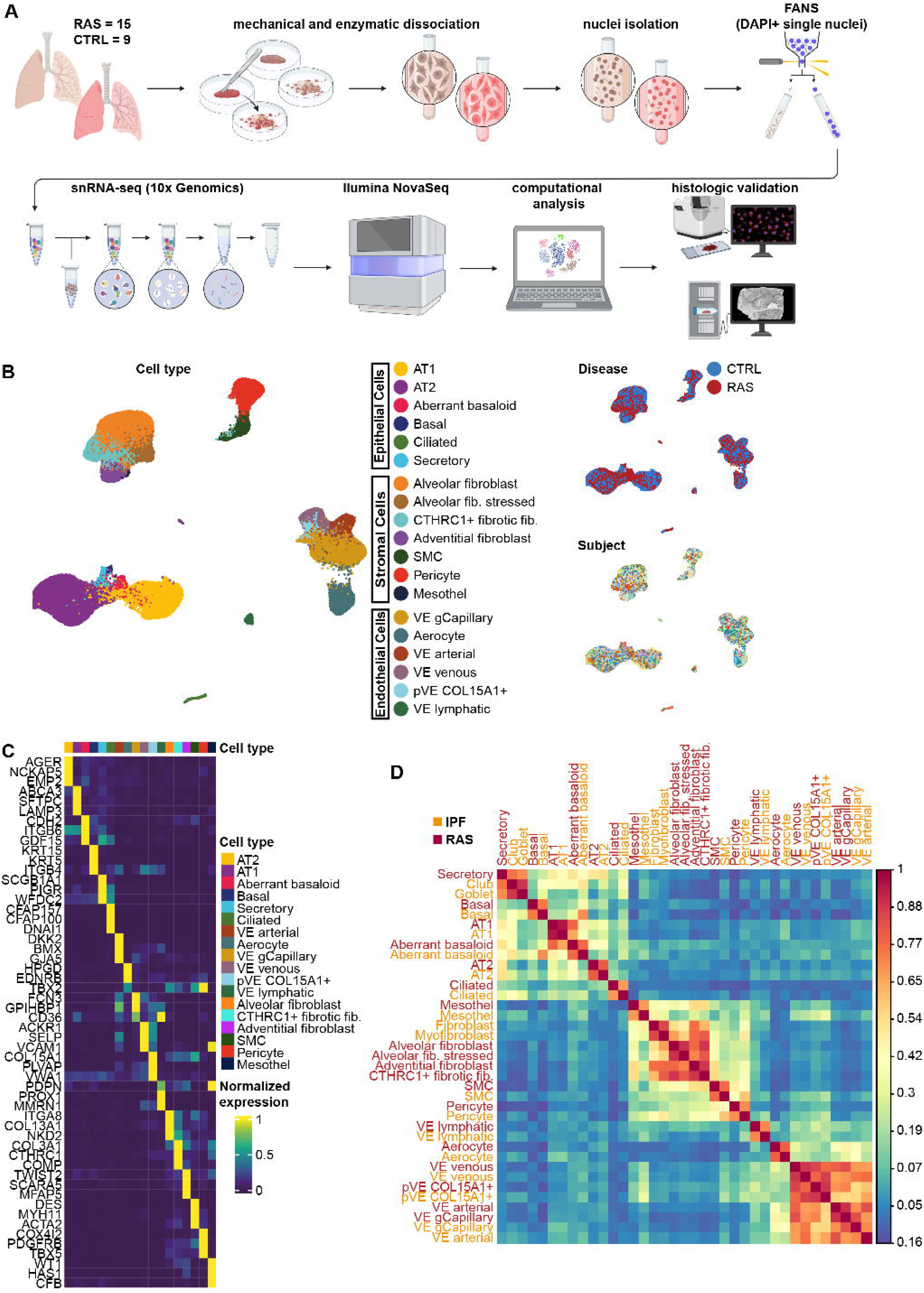
**A**: Overview of experimental design. 1) Tissue samples of 15 RAS lung explants from patients undergoing lung re-transplantation and from 9 donor control lungs were used. 2) + 3) Lung tissue samples were mechanically and enzymatically dissociated. 4) Nuclei were isolated from dissociated lung tissue specimen. 5) Flow activated nuclei sorting (FANS) using DAPI was performed to extract a clean single nuclei suspension of each specimen. 6) SnRNA-seq library preparation was performed using 10x Genomics’ Chromium Next GEM technique for multiplexed fixed samples. 7) Sequencing of snRNA-seq libraries using the Ilumina platform. 7) Computational explorative analysis and validation of snRNA-seq data. 8) Histologic validation and spatial localization by immunofluorescence, RNAscope, combined Elastica van Gieson-immunohistochemistry stains, and micro-CT scans. **B**: Uniform Manifold Approximation and Projection (UMAP) representation of 105,952 single nuclei obtained from 15 RAS, and 9 control donor lungs. Each dot represents a single nucleus. Nuclei are labeled as one of 19 distinctive cell types in the left UMAP: epithelial cells (AT1, AT2, aberrant basaloid, basal, ciliated and secretory cells), endothelial cells (aerocytes, vascular endothelial (VE) arterial, VE gCapillary, VE lymphatic, VE systemic, and VE venous cells), and stromal cells (Fibroblasts (alveolar, adventitial, fibrotic, and inflamed alveolar), mesothel, pericytes, and smooth muscle cells (SMC)). Additionally, nuclei are labeled by disease status (top right), and subject (bottom right) where each color depicts a distinct subject. **C**: Heat map of the average marker gene expression per subject per cell type. Gene expression values are unity normalized from 0 to 1. **D**: Correlation matrix of all identified cell populations of this dataset and of analogous parenchymal cell types of the IPF atlas(13). Matrix cells are colored by Spearman’s rho. Annotation bars (RAS or IPF) denote the origin dataset of each cell population.

## MATERIAL AND METHODS

Complete experimental details are provided in the Methods section in the Supplement. Briefly, we performed single-nuclei RNA-sequencing (snRNA-seq) using peripheral lung parenchyma from upper lobes of 15 RAS patients undergoing lung ReTx which were phenotyped according to the latest ISHLT criteria and of 9 control lungs (surgigal size adjustement during lung transplantation n=5; histologically tumor-free specimen n=4); Fig. 1A).(5) SnRNA-seq data were processed with CellRanger and analyzed using R and the Seurat package. Epithelial, endothelial and stromal cell populations were characterized through iterative clustering followed by differential expression analysis. Wilcoxon rank sum test with *P* values adjusted for multiple comparisons using the Bonferroni method established cell type-specific marker genes. Epithelial, endothelial and stromal cell subpopulations were validated and localized by immunohistofluorescent (IHF) microscopy, RNAscope, and combined Elastica van Gieson-immunohistochemistry (EvG-IHC) stains. Micro-CT scans of our specimens provided further insights in the structural characteristics of RAS. This study was approved by the Ethics Committee of Hanover Medical School (IRB # 10141_BO_K_2022).

## RESULTS

SnRNA-seq data of 105,952 single nuclei was analyzed, of which 15 originate from RAS specimens and 9 from control specimens (Fig. 1B). A technical summary including median UMI/genes/reads per nuclei, reads mapped to probe set, and the number of detected genes is provided in the supplement (Supp. Table S2A). Based on distinct markers, all major cell types of the human lung were identified, in addition to aberrant epithelial, an ectopic endothelial and a fibrotic fibroblast population that are idiopathic pulmonary fibrosis (IPF) -associated and were previously undiscovered in RAS (Fig. 1B-D). The detailed marker gene expressions of all profiled nuclei are provided in the supplement (Supp. Table S2B).

### SnRNA-seq analysis of the epithelial cell repertoire reveals aberrant basaloid cells in human RAS lungs

40,390 single nuclei representing 38.1% of all profiled single nuclei were identified as epithelial cells based on distinct marker genes. We identified most known major epithelial cell populations of the human lung, including alveolar type 1 (AT1) and type 2 (AT2) cells, basal cells, ciliated cells, and secretory cells in all specimens (Figs. 2A-D; Supp. Table S2C). In addition, we could identify the presence of aberrant basaloid cells in all but one of our RAS specimens. This characteristic disease-enriched cell type was first described in 2020 by Adams et al. and Habermann et al. in human IPF lungs, and to a lesser extent in COPD lungs.(12,13) Our data reveals a distinctive gene signature of these aberrant basaloid cells that aligns with the description of Adams et al.(13): Aberrant basaloid cells express the basal cell markers KRT17, and LAMB3 in addition to canonical epithelial markers, but lack the expression of other established basal cell markers such as KRT5 and KRT15 (Fig. 2D). Furthermore, aberrant basaloid cells express senescence-related genes like CCND1, MDM2, and GDF15 as well as markers of epithelial-mesenchymal transition (EMT) such as CDH2, COL1A1, VIM, and FN1.(13) They are characterized by the highest expression levels of genes coding for the integrin subunits αvβ6, activators of TGFβ, compared to all other cell types of our specimens. We identified 1,274 aberrant basaloid cells matching this unique gene signature, representing in the median 5.4% of the epithelial cell compartment in RAS patients (vs. controls, p < 0.0001, Wilcoxon rank-sum test, Fig. 2B, 2C; Supp. Table S2H). Due to the integration approach of our snRNA-seq data analysis, median 0.7% nuclei of our control specimens were identified as aberrant basaloid cells, too. However, analysis of their gene expression profile revealed a substantial reduced expression of major aberrant basaloid-specific marker genes (Fig. 2D). Correlation analysis with the IPF cell atlas data confirmed the transcriptional concordance of aberrant basaloid cells in RAS and IPF (Fig. 1D).(13)

**Figure 2.**
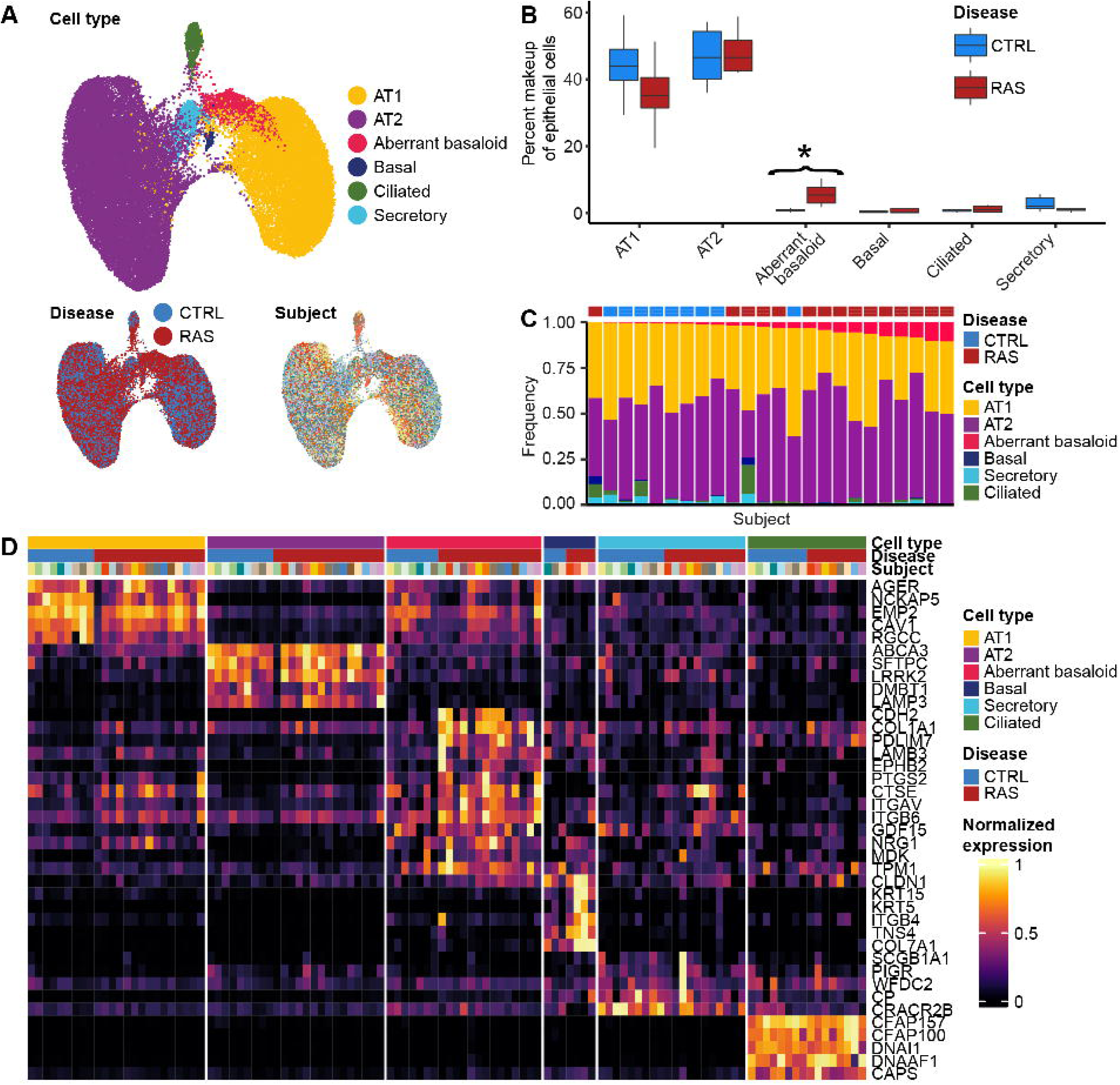
**A**: UMAPs of 40,390 epithelial single nuclei from 15 RAS, and 9 control donor lungs, labeled by cell type (upper UMAP), disease (bottom left), and subject (bottom right) where each color depicts a distinct subject. **B**: Boxplots representing the percent makeup distributions of each epithelial cell type among all identified epithelial cells organized by disease group. Whiskers represent 1.5 × interquartile range (IQR). Results of FDR-adjusted Wilcoxon rank sum test comparing RAS and control proportions are reported in supplemental table S2H. **C**: Stacked bar plots representing the frequency (in %) of each epithelial cell type among all identified epithelial cells per subject. Each bar represents a different subject (15 RAS and 9 control subjects). Stacked bars are ordered by increasing frequencies of aberrant basaloid cells. **D**: Heat map representing characteristics of the six identified epithelial cell types. Gene expression is unity normalized between 0 and 1 across epithelial cells. Each column shows the average expression per subject and disease state. Each subject is represented by a unique color, and disease state and epithelial cell type are represented in the colored annotation bars above.

### Aberrant basaloid cells are uniquely localized at the edge of the fibrotic pushing border

We confirmed the presence of aberrant basaloid cells in RAS tissue and investigated their distribution by IHF using antibodies against KRT17, TP63, CTSE, integrin αvβ6, and COX2, which represent a combination of distinctive markers that does not overlap with any other common cell type of the human lung (Fig. 2D).(13) All IHF stainings of our RAS samples showed the same characteristic distribution pattern of aberrant basaloid cells: They are consistently located at the edge of the fibrotic pushing border, and line it almost continuously in a single cell layer (Fig. 3). In addition, aberrant basaloid cells reach foothill-like into the alveolar space, where they line fibrotically thickened alveolar septa (Fig. 3). Furthermore, we throughout observed an epithelial gradient in RAS: Aberrant basaloid cells occur with high frequencies at the fibrotic pushing border and become increasingly rare as they extend into areas of presumably healthy-looking alveoli until they eventually disappear. Interestingly, all aberrant basaloid cells are CTSE and integrin αvβ6 positive, but CTSE+/αvβ6+ cells themselves reach further into presumably healthy-looking alveoli and are more frequently than aberrant basaloid (Fig. 3). In contrast to IPF, myofibroblast foci covered by aberrant basaloid cells are a rather rare histological finding in RAS. In control specimens, we could not detect any aberrant basaloid cells.

**Figure 3.**
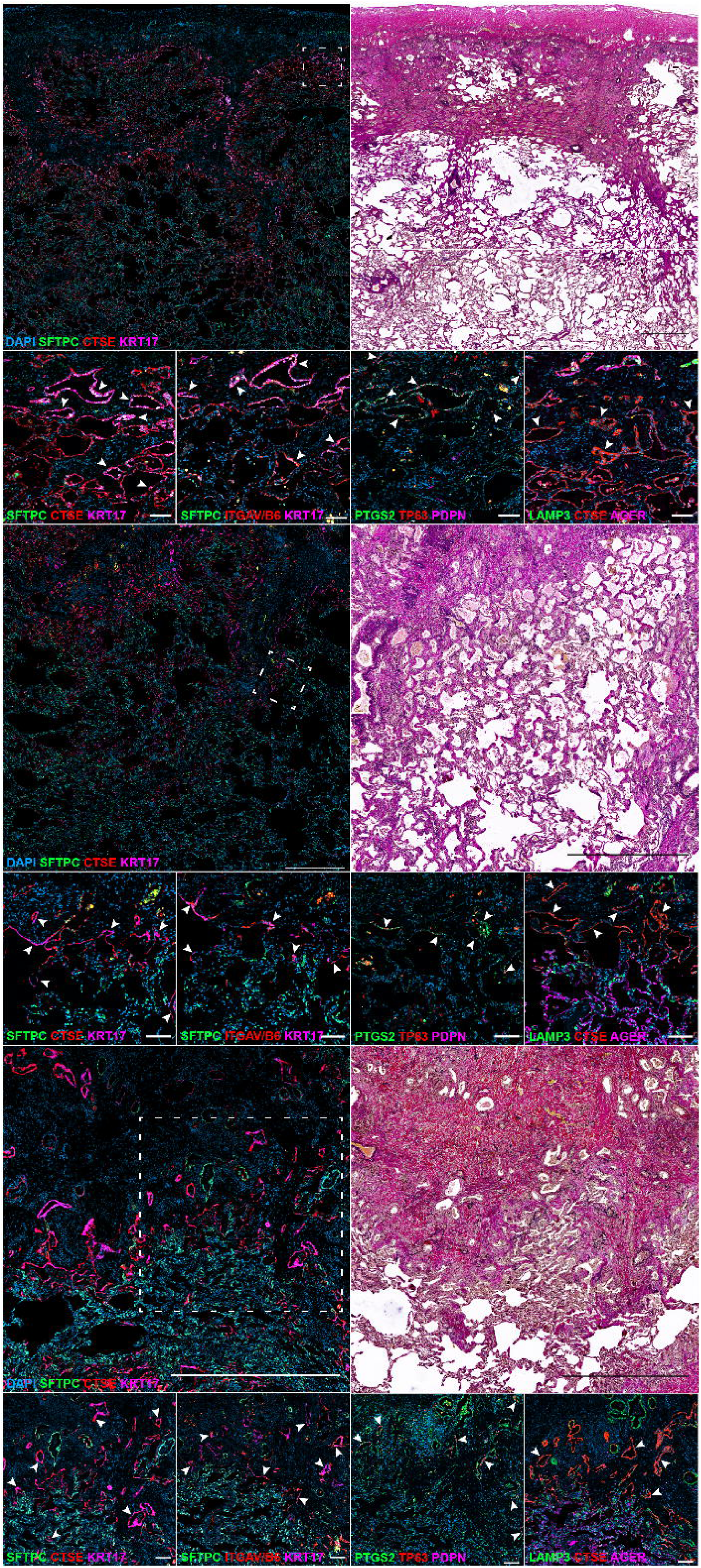
IHF and EvG stains of RAS specimens. Scale bar of overviews (row 1, 3, and 5) are 1000 µm. Scale bars of close-up views (rows 2, 4, and 6) are 100 µm. Arrows indicate aberrant basaloid cells. Control stains are depicted in supplemental figure Fig. S2.

Fully developed AFE is characterized by a homogeneous, pauci-cellular fibrosis with a preservation of the elastic fibers of the remnant alveolar walls. These are denuded of epithelial cells, while the former alveoli are drowned in collagen. However, in proximity to the fibrotic pushing border, we consistently observed multiple air-containing structures, which we termed “obliterating alveoli”. Lining epithelial cells of obliterating alveoli have lost contact to remnant alveolar walls due to thick collagen depositions. Obliterating alveoli consist of either aberrant basaloid cells, SFTPC+/KRT17+ transitional cells, and LAMP+/AGER+ AT1 cells, all of which being pathologically CTSE+ (Fig. 3). Even though any combination between these cell types exists, obliterating alveoli are predominantly formed by aberrant basaloid cells, and most heterogenous obliterating alveoli contain at least some aberrant basaloid cells. In general, aberrant basaloid seem to exclusively appear in monolayers, independent of their localization in RAS lung tissue.

### The endothelial cell repertoire is shifted towards ectopic COL15A1+ vascular endothelial cells, and shows a loss of normal alveolar microvasculature

27,657 single nuclei representing 26.1% of all profiled single nuclei were identified as endothelial cells (ECs) based on distinct markers (Fig. 4A-D). We identified all known EC populations of the human lung in all specimens, including arterial ECs, aerocyte, general capillary ECs, venous ECs, lympathic ECs, and COL15A1+ systemic ECs (Supp. Table S2D). Since this last vascular EC (VEC) population transcriptomically matches the description of peribronchial VE (pVE) cells, which form vessels adjacent to major airways and subpleural vessels, we strictly refer to them as COL15A1+ pVE cells (Fig 4D).(13) The endothelial repertoire in RAS is shifted to a significantly increased fraction of COL15A1+ pVE cells (1.9% vs. 8.2%, p < 0.05, Wilcoxon rank-sum test, Fig. 4B; Supp. Table S2H). Contrasting, we observe a non-significant trend towards a decreased fraction of aerocytes in RAS (median percentage 26.0% vs. 11.2%, p = 0.055; Fig. 4B).

**Figure 4.**
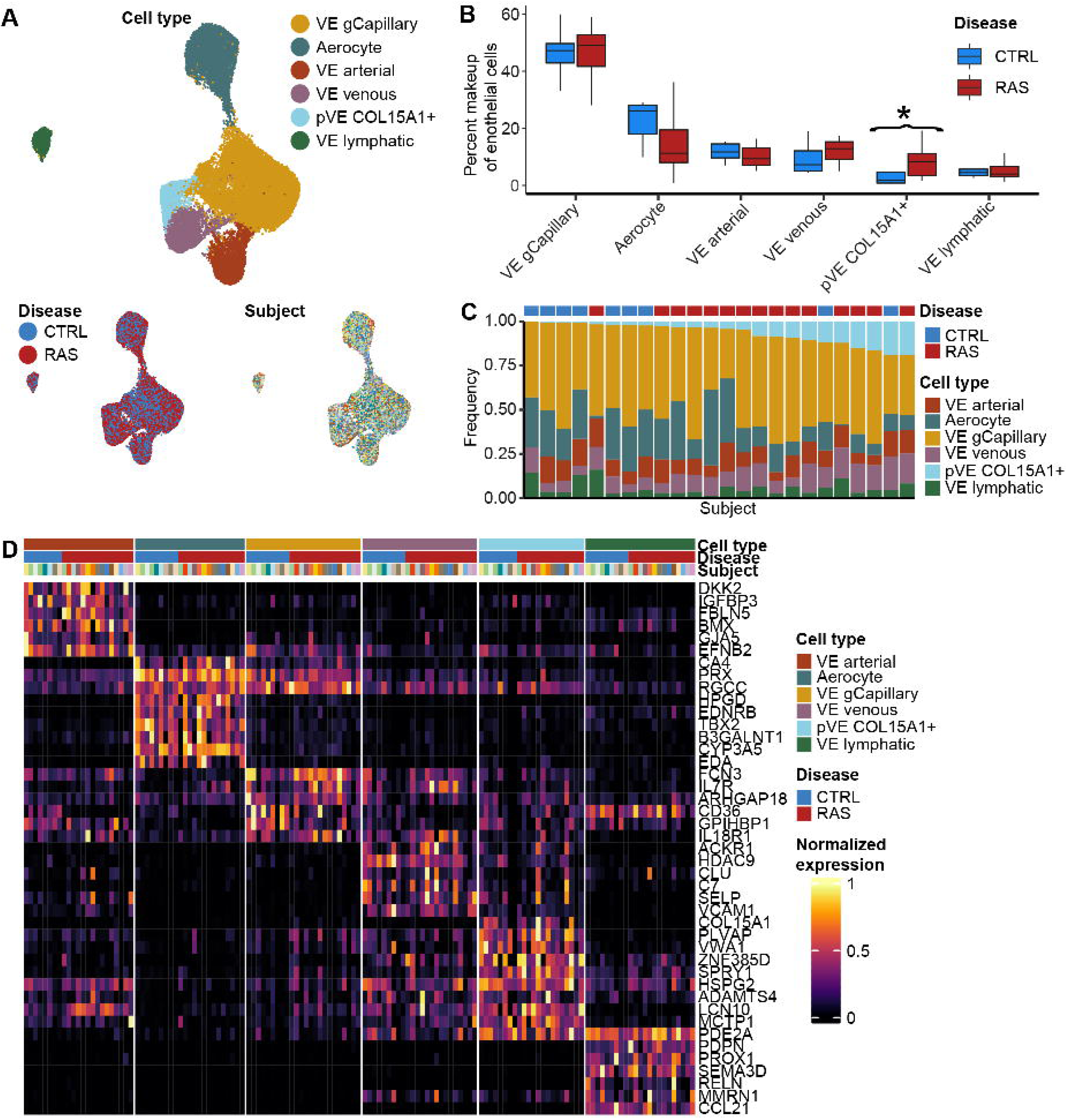
**A**: UMAPs of 27,657 epithelial single nuclei from 15 RAS, and 9 control donor lungs, labeled by cell type (upper UMAP), disease (bottom left), and subject (bottom right) where each color depicts a distinct subject. **B**: Boxplots representing the percent makeup distributions of each endothelial cell type among all identified endothelial cells organized by disease group. Whiskers represent 1.5 × interquartile range (IQR). Results of FDR-adjusted Wilcoxon rank sum test comparing RAS and control proportions are reported in supplemental table S2H. **C**: Stacked bar plots representing the frequency (in %) of each endothelial cell type among all identified endothelial cells per subject. Each bar represents a different subject (15 RAS and 9 control subjects). Stacked bars are ordered by increasing frequencies of COL15A1+ systemic VEs. **D**: Heat map representing marker gene expression of the six identified endothelial cell types. Gene expression is unity normalized between 0 and 1 across endothelial cells. Each column shows individual cell average marker gene expression per subject and disease state. Each subject is represented by a unique color, and disease state and endothelial cell type are represented in the colored annotation bars above.

### IHF stains reveals ectopic pVE cells in RAS, which partially replace normal PRX+ alveolar microvasculature

We validated the presence of pVE cells using antibodies against their distinctive markers COL15A1 and PLVAP alongside the pan-endothelial marker CD31. In control specimens, CD31+/COL15A1+ pVE cells are strictly restricted to peribronchial vasculature or to larger vessels of the pleura, but do not contribute to the normal PRX+ alveolar microvasculature (Supp. Fig. S2D). In RAS, COL15A1+ pVE cells show a deviating distribution pattern: CD31+/COL15A1+ vessels are no longer restricted to peribronchial vasculature, but expand dramatically (Fig. 5A). We refer to all COL15A1+ pVE cells apart from locations adjacent to major airways or the pleura as ‘ectopic pVE cells’. Remarkably, ectopic pVE cells gradually expand parallel running to the aberrant basaloid cell gradient: They are primarily located in vessels of varying sizes in fibrotic areas adjacent to the fibrotic pushing border (Fig. 5A). Towards the dense fibrotic subpleural space, we observe a rarefication of ectopic pVE cells, but an increase in the diameter of the respective vessels. Similar to our epithelial findings, ectopic pVE cells are not restricted to fibrotic areas, but invade alveolar tissue starting from the fibrotic pushing border. Where ectopic pVE cells reach into alveolar tissue in shape of microvasculature, we observe a loss of physiologic CD31+/PRX+ alveolar microvasculature in both, our IHF and EvG-IHC stainings (Fig. 5A, 5B).

**Figure 5.**
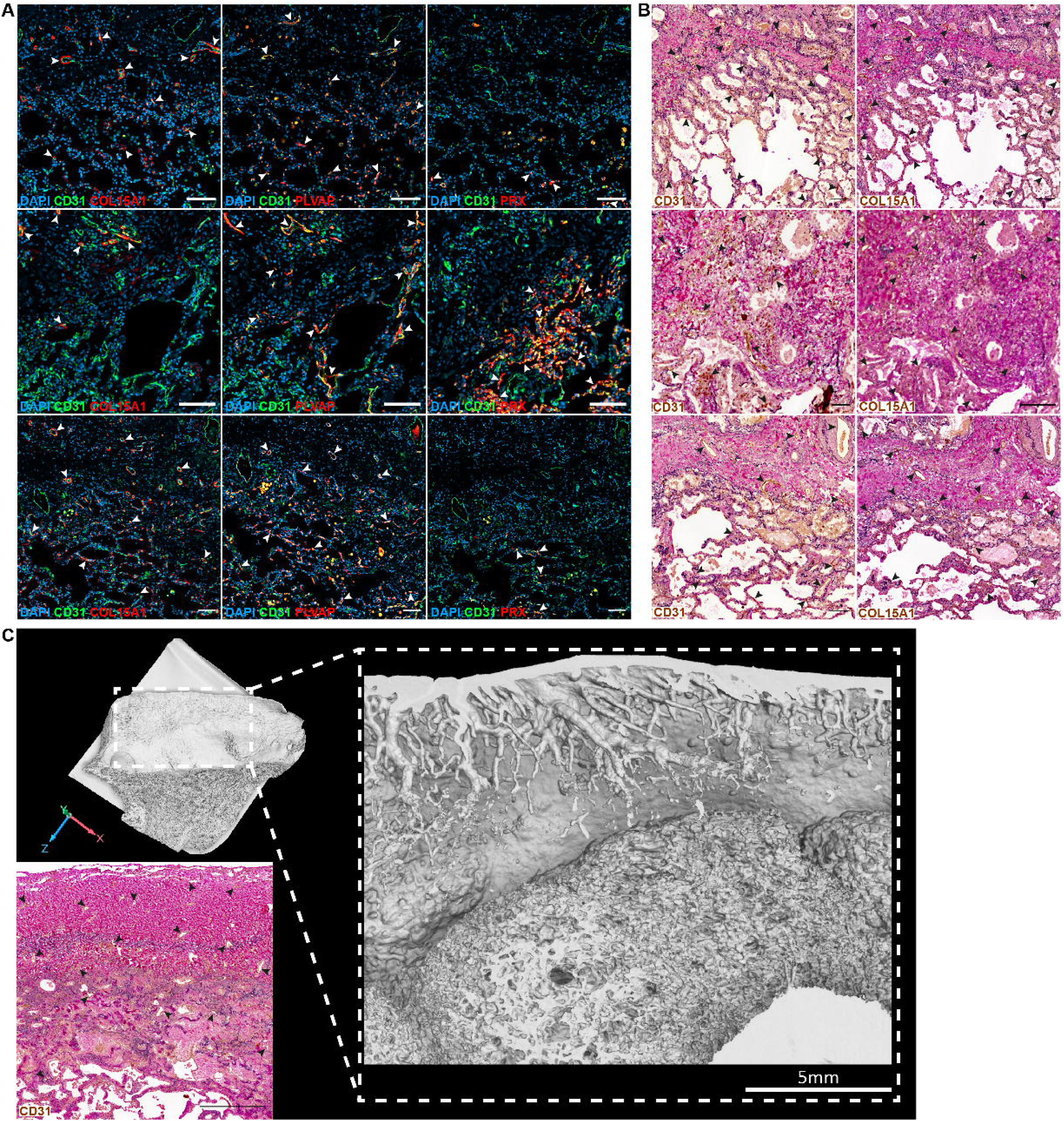
**A**: IHF stains of RAS specimens. Scale bars are 100 µm. Arrows A (insert arrow) indicate *COL15A1+/PLVAP+* ectopic peribronchial VE cells. Arrow B (insert arrow) indicates *PRX+* microvasculature. **B**: IHC-EvG stainings of RAS specimens. Scale bars are 100 µm. Arrows indicate *CD31+* VE cells and *COL15A1+* ectopic peribronchial VE cells. **C**: Micro-CT imaging of subpleural RAS remodeling. RAS tissue specimen was scanned with a Phoenix Nanotom® M micro-CT scanner (Waygate technologies) following de-paraffinization and contrasting with Tungsten phosphoric acid for 48h. 3D reconstruction of the tissue specimen shows a dense zone of subpleural fibrotic consolidation. Tungsten contrasting does not contrast fibrotic tissue. Virtual sectioning of the block indicates pronounced sprouting of vasculature-like structures within the subpleural fibrosis. Morphologically, vascular structures seem to originate from the pleura, as cross sections of the observed vessels rejuvenate with increasing distance from the pleura. IHC-EvG staining of a RAS specimen. Scale bar is 500 µm. Arrows indicate *CD31+* VE cells. Control stains are depicted in supplemental figure Fig. S3.

Micro-CT imaging with 3D reconstruction reveal pronounced vascular remodeling in the subpleural space in the dense zone of subpleural fibrotic consolidation in RAS tissue (Fig. 5C). Virtual sectioning indicates pronounced sprouting of multiple vasculature-like structures within the subpleural fibrosis reaching into AFE areas. Sequential EvG-IHC stainings confirm endothelial lining vasculature-like structures due to positive CD31 staining (Fig. 5B). Morphologically, vascular structures seem to originate from the pleura, as cross sections of the observed vessels rejuvenate with increasing distance from the pleura (Fig. 5C).

### The stromal cell compartment is shifted towards *CTHRC1+* fibrotic fibroblasts in RAS

37,905 single nuclei representing 35.8% of all profiled single nuclei were identified as stromal cells based on distinct markers (Fig. 6A-D, Supp. Table S2E). They include pericytes, smooth muscle cells (SMC), mesothelium, and fibroblast populations. Within fibroblasts, we identified distinctive subpopulations, including alveolar fibroblasts, inflamed alveolar fibroblasts, adventitial fibroblasts, and *CTHRC1+* fibrotic fibroblasts. While *CTHRC1+* fibrotic fibroblasts are found in RAS and control subjects, they compose a significantly higher fraction of all stromal cells in RAS compared to controls (median percentage among stromal cells: 21.5% RAS vs. 6.4% control, p = 0.0002; Fig. 6B; Supp. Table S2H). In contrast, the fraction of alveolar fibroblasts in RAS are significantly reduced (RAS vs. control: 38.0% vs. 48.5%, p = 0.005; Fig. 6B). The transcriptional profile of *CTHRC1+* fibrotic fibroblasts in our snRNA-seq data matches published descriptions of the IPF-associated *CTHRC1+* myofibroblast subpopulation (13,14): Aside from CTHRC1, they exhibit the highest expression of collagens COL1A1 and COL3A1, ECM-related genes such as COMP, LUM, SPARC, THBS2, POSTN, FN1, the growth factor VEGFA, the matricellular protein CCN4, the TGF-β signaling pathway activator INHBA, and the secreted serine protease PLAU (Fig. 6D, 6E). Further, they share common gene markers of alveolar fibroblasts such as including PDGFRA, TCF21, and NPNT (Fig. 6D).(14) Biological pathways enriched in CTHRC1+ fibrotic fibroblasts include: elastic fiber formation; chondroitin sulfate, dermatan sulfate and peptide hormone biosynthesis; activation of matrix metalloproteinases; collagen degradation, collagen chain biosynthesis and modifying enzymes, and collagen chain trimerization (Fig. 6F; Supp. Table S2F). Taken together, marker gene expression and enriched pathways of CTHRC1+ fibrotic fibroblasts highlight their prominent role in the fibrotic remodeling of AFE.

**Figure 6.**
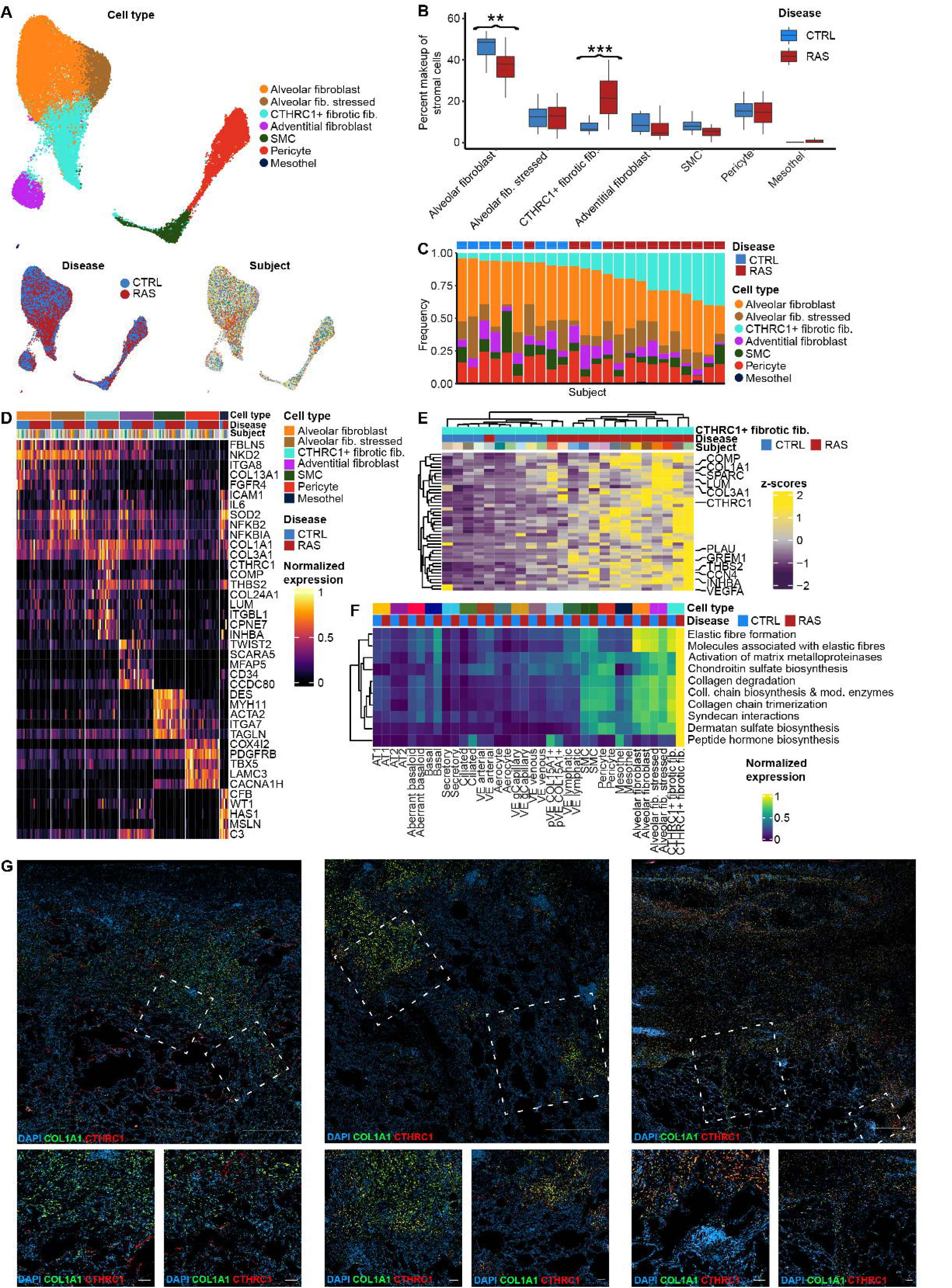
**A:** UMAPs of 37,905 stromal single nuclei from 15 RAS, and 9 control donor lungs, labeled by cell type (upper UMAP), disease (bottom left), and subject (bottom right) where each color depicts a distinct subject. **B:** Boxplots representing the nonzero percent makeup distributions of each stromal cell type among all identified stromal cells organized by disease group. Whiskers represent 1.5 × interquartile range (IQR). Results of FDR-adjusted Wilcoxon rank sum test comparing RAS and control proportions are reported in supplemental table S2H. **C**: Stacked bar plots representing the frequency (in %) of each stromal cell type among all identified stromal cells per subject. Each bar represents a different subject (15 RAS and 9 control subjects). Stacked bars are ordered by increasing frequencies of CTHRC1+ fibrotic fibroblasts. **D**: Heat map representing the marker gene expression of the seven identified stromal cell types. Gene expression is unity normalized between 0 and 1 across stromal cells. Each column shows average gene expression per subject and disease state. Each subject is represented by a unique color, and disease state and stromal cell type are represented in the colored annotation bars above. **E**: Heat map of differentially expressed genes comparing RAS and Controls in CTHRC1+ fibrotic fibroblasts, hierarchically clustered. Average expression values are scaled across subjects. Each subject is represented by a unique color, and disease state and cell type are represented in the colored annotation bars above. **F**: Pathway analysis of 15 RAS and 9 control specimens. Depicted are average enrichment scores per cell type and disease state. Enrichment scores are unity normalized between 0 and 1 across all identified cells types. Cell type and disease status are indicated by the colored annotation bars above. **G**: RNAscope stainings of RAS specimens. Scale bar of overviews (top row) are 1000 µm. Scale bars of close-up views (bottom row) are 100 µm. Arrows indicate CTHRC1+ fibrotic fibroblasts.

**Figure 7.**
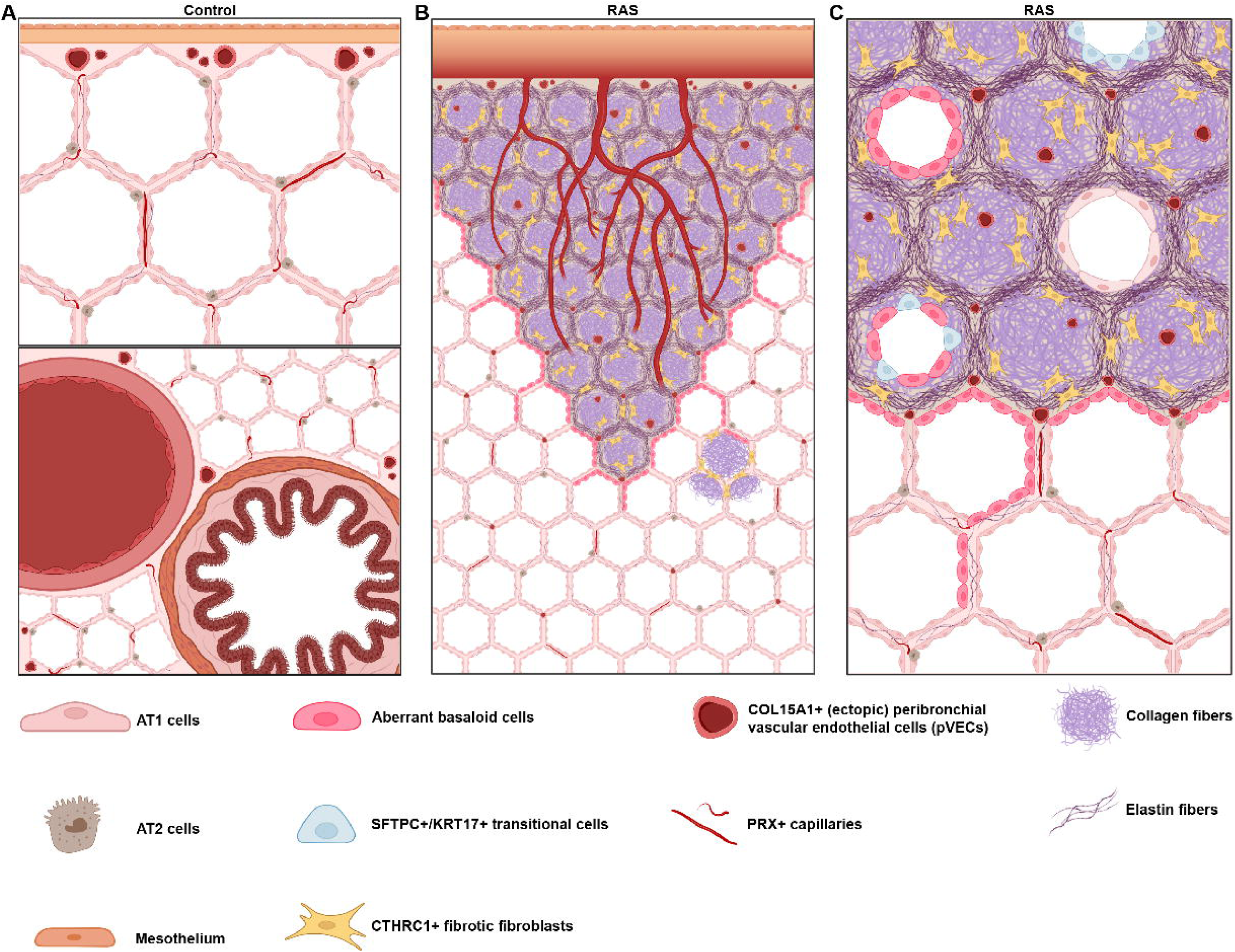
**A:** Schematic illustration of healthy lung tissue. Upper illustration shows the mesothelial coated pleura, and subpleural lung tissue including *COL15A1+* vessels near the pleura, alveoli, elastin fibers in alveolar septa, AT1 and AT2 cells, and *PRX+* capillaries. Bottom illustration shows healthy lung tissue including a bronchus and its corresponding bronchial artery, peribronchial *COL15A1+* vessels, alveoli, elastin fibers in alveolar septa, AT1 and AT2 cells, and *PRX+* capillaries. **B:** Schematic overview illustration of RAS lung tissue. It shows characteristic elements of RAS such as a thickened pleura, areas of alveolar fibroelastosis (AFE), vasculature-like structures within the subpleural fibrosis reaching into AFE areas and rejuvenating with increasing distance from the pleura, *COL15A1+* vessels near the pleura and ectopic *COL15A1+* vessels in areas of AFE and in alveolar septa where they partially replace *PRX+* capillaries, a decreased amount of *PRX+* capillaries, *CTHRC1+* fibrotic fibroblasts in areas of AFE and sporadically in small nests in thickened alveolar septa in more distal RAS tissue, aberrant basaloid cells lining the active edge of the fibrotic pushing border in a monolayer and lining thickened alveolar septa, and enriched elastic fibers in thickened alveolar septa. **C:** Schematic close-up illustration of RAS lung tissue. It shows characteristic elements of RAS such as AFE, ectopic *COL15A1+* vessels in areas of AFE and in alveolar septa where they partially replace *PRX+* capillaries, a decreased amount of *PRX+* capillaries, *CTHRC1+* fibrotic fibroblasts in areas of AFE, aberrant basaloid cells lining the active edge of the fibrotic pushing border in a monolayer and lining thickened alveolar septa, obliterating alveoli near the fibrotic pushing border consisting of either aberrant basaloid cells, SFTPC+/KRT17+ transitional cells, and LAMP+/AGER+ AT1 cells, all of which being pathologically CTSE+, and enriched elastic fibers in thickened alveolar septa.

### RNAscope reveals localization of *CTHRC1+* fibrotic fibroblasts in RAS

RNAscope stains of *CTHRC1+/COL1A1+* fibrotic fibroblasts reveal a predominant localization of *CTHRC1+* fibrotic fibroblasts in areas of AFE in RAS, and only very few *CTHRC1+* fibrotic fibroblasts were observed in controls (Fig. 6G). More specifically, *CTHRC1+* fibrotic fibroblasts are ubiquitously distributed across all areas of AFE, partially forming very few fibroblast foci, especially at the fibrotic pushing border. In contrast to aberrant basaloid and ectopic pVE cells, *CTHRC1+* fibrotic fibroblasts are not specifically enriched at the fibrotic pushing border. Apart from fibrotic AFE areas, *CTHRC1+* fibrotic fibroblasts were sporadically observed in small nests in thickened alveolar septa in more distal RAS tissue, but not in presumably physiologically-looking, thin alveoli walls (Fig. 6G).

## DISCUSSION

In this study, we provide the first-ever cellularly-resolved transcriptomic data of human RAS lung tissue. Our snRNA-seq data reveals the presence of aberrant basaloid cells in the epithelial compartment, and a shift towards CD31+/COL15A1+ vascular endothelial cells, which overlap with the description of physiologic pVE cells and ectopic pVE cells by Adams et al.(13) Our IHF stains confirm the existence of aberrant basaloid cells, and ectopic pVE cells in RAS and their absence in control lungs. In addition, specific and unique distribution patterns of both cell populations were revealed by our IHF, EvG and HE stains. The existence of aberrant basaloid cells, ectopic pVE cells, and *CTHRC1+* fibrotic fibroblasts was first described in idiopathic pulmonary fibrosis (IPF), then confirmed in patients with lung involvement of scleroderma(15), but was previously unknown in RAS lung tissue.

The role and contribution of parenchymal cell in general, and epithelial cells specifically, in the pathogenesis of RAS has recently shifted more into focus.(7) Here, add to it the surprising discovery of rare and unique aberrant basaloid cells among the epithelial cell repertoire in RAS. Aberrant basaloid cells are disease-associated cells, which have been found in IPF and scleroderma with pulmonary involvement (SSc-ILD), and to a lesser extent in COPD.(12,13,15) The origin of aberrant basaloid cells remains elusive, but latest research indicates that human AT2 cells are able to transdifferentiate into aberrant basaloid cells.(16) Their detection in the human RAS lung implies previously unknown commonalities of these clinically completely different entities with potential impact on therapeutic targets. Our expression profile of these cells completely matches the aberrant basaloid cell population detected by Adams et al.(13) in human IPF lungs despite deviating distribution patterns: in IPF, aberrant basaloid cells characteristically line the active edge of pathognomonic myofibroblast foci.(13) In contrast to IPF, fibroblast foci are rare in the RAS lung. However, a nearly continuous monolayer of aberrant basaloid cells is formed at the active edge of the fibrotic pushing border. Combined with their extended penetration covering fibrotically thickened alveolar walls, our findings implicate a more general involvement of these cells in the pathogenesis of RAS. In contrast to IPF, bronchiolization is absent in RAS, and it was neither observed in our histological stains nor in terms of frequencies of bronchial cell types in our snRNA-seq data.

Our data indicates a relevant influence of this cell population on the pathogenesis and/or disease progression of RAS: firstly, we identified aberrant basaloid cells in all but one of our RAS specimens, but no typical aberrant basaloid cell in control specimens. Consequently, our data supports the hypothesis of aberrant basaloid cells as a pathognomonic feature of fibrotic lung diseases in general, and as a hallmark feature of RAS specifically. Secondly, their unique expression profile may promote the fibrotic remodelling seen in regions of AFE in RAS lungs. This is indicated by the epithelial mesenchymal transition-like phenotype (EMT) due to the expression of CDH2, COL1A1 and VIM, and the uniquely high expression of genes associated with the pathogenesis of IPF such as integrin αVβ6, MMP7, GDF15, and EPHB2. Thirdly, cellular senescence is associated with disease progression i.e., of age-related and chronic diseases like IPF as well as frailty and transplant-specific complications in solid-organ transplants.(17) Hence, it is likely that aberrant basaloid cells reflect drivers of pathologic cellular senescence in RAS due to their expression of multiple senescence markers like CDKN1A, CDKN2B, MDM2, and GDF15. Fourthly and most importantly, our data shows strongly elevated expression levels of genes coding for the integrin subunits αvβ6 in aberrant basaloid cells. Integrin αvβ6 can activate latent TGF-β1, and high expression levels of integrin αvβ6 are known to substantially contribute to organ fibrosis.(18) Increased expression levels of TGF-β1 were already reported in BALF analysis from RAS patients and associated with the development of BOS by Morrone and colleagues.(19,20) Integrin αvβ6 is therefore considered as a promising potential therapeutic target for IPF. Our findings now strongly suggest considering integrin αvβ6 as a potential novel therapeutic molecular target in RAS as well, and provide rationale for transferability of future aberrant basaloid and epithelial cell-targeted IPF therapeutics to RAS due to the diseases‘ transcriptomic similarities.

To date, little is known about the endothelial cell compartment in RAS patients and its possible pathologic impacts. As in IPF, we observed an significant expansion of COL15A1+ ectopic pVE cells in RAS, with largely overlapping transcriptomic profile.(13) Physiologically, pVE cells are restricted to the pleura and the vascular plexus surrounding larger airways and vessels, but are transcriptomically indistinguishable from ectopic pVE cells in RAS.(21) Histo-anatomically, the distribution of ectopic pVE cells differs between IPF and RAS in some aspects: In IPF, CD31+/COL15A1+ ectopic pVE cells are associated with areas of bronchiolization, and therefore fit the cellular proximalization of distal IPF lung tissue. In RAS, our data confirms the absence of bronchiolization. Instead, ectopic pVE cells at the fibrovascular interface are characteristically distributed parallel-running to the aberrant basaloid cell gradient.(22) We often observed a rarefication of these vessels towards the center of fibrotic areas, but high frequencies in proximity to the fibrotic pushing border. Toward the pleura, the vascular diameter markedly increases, as observed histologically and in micro-CT scan, suggestive for a change from pulmonary to systemic circulation in areas if AFE. Interestingly, wherever ectopic pVE cells infiltrate the alveolar microvasculature, CD31+/PRX+ physiologic alveolar capillaries are missing. Although our data cannot determine the function of capillary-like ectopic pVE vessels, we doubt a complete physiologic capacity. Physiologically, the lung microvasculature is of the continuous type. In RAS, PLVAP - specifically expressed in ectopic pVE cells - is known to form the diaphragms of fenestrated endothelium, to substantially influence vascular permeability and leukocyte trafficking/transmigration and is only pathologically expressed in mature barrier endothelium.(23,24) Therefore, we assume ectopic pVE cells substantially impact i.e., gas exchange, vascular permeability, and prostaglandin homeostasis in the RAS lung.

The origin and role of ectopic pVE cells in RAS remains enigmatic nonetheless. Our data cannot determine if the described gradients of aberrant basaloid cells and ectopic pVE cells are connected. However, the specific location of these parallel-running gradients, and the highest frequencies of both cell populations at the fibrotic pushing border imply them as disease-specific features, potentially due to repetitive endothelial injury.

Fibroelastotic remodeling in the areas of the remnant alveolar walls is the hallmark pathological feature in RAS lung tissue.(10,25–27) One cell population stood as cellular mediator of this fibroelastotic remodeling in RAS: CTHRC1+ fibrotic fibroblasts(13,14), as they express numerous genes associated with fibrosis including BMP1, COL1A2, COL3A1, CXCL12, CTHRC1, MMP2, MMP14, PLOD2, SERPINE1, TIMP1, and TIMP2.(14) Furthermore, pathway analysis in CTHRC1+ fibrotic fibroblasts revealed characteristic pathomechanisms of fibroelastotic remodeling, including elastogenesis, collagenesis, elastic fiber and ECM formation, organization, and its regulation. The fibroblast to CTHRC1+ myofibroblast conversion through SFRP1+ transitional fibroblasts has been recently unravelled.(28) CTHRC1+ fibrotic fibroblasts have been identified as key driver of fibrosis, not only in the lung as in IPF and scleroderma-ILD, but in organ fibrosis throughout.(29–31) Accordingly, our study revealed CTHRC1+ fibrotic fibroblasts as key effector of fibroelastotic remodeling in RAS as well, adding to their prominent role in organ fibrosis in general.

Our study is not without limitation. First, the size of our cohort does not allow capturing the complete heterogeneity of RAS, although analysis of 15 RAS patients is a decent cohort size for such a rare disease. Second, analyzing end-stage RAS tissue provides insights into the pathophysiologic characteristics of RAS, which might be more ambiguous or lacking in earlier disease stages. Third, because we provide nuclei data, it is principally not possible to capture the entirety of genes in our data, but we still cover the large majority of genes on a cellularly-resolved base. In future, it will be necessary to generate further single-cell or single-nuclei RNAseq data sets in order to confirm and complement our findings.

Taken together, this study - together with previous scRNAseq studies in fibrotic lung diseases(12,13,15) - reveals a striking, but common cellular similarity in fibrotic lung diseases, despite impressive histo-morphological differences, such as the UIP pattern in IPF and AFE pattern in RAS: We observed that the fibrotic niche is likewise formed by aberrant basaloid cells, ectopic *COL15A1+* pVE cells, and *CTHRC1+* fibrotic fibroblasts. Therefore, our study offers strong arguments investigating the transferability of recent and future IPF-specific therapeutics to a RAS-specific therapy, since the unexpected transcriptomic similarity of disease-relevant cell types in IPF and RAS has not been clinically anticipated.

## ACKNOWLEDGMENTS

We are indebted with great gratitude to all patients who enabled conducting this study by their participation. We thank AG Fiedler, AG Braun, and AG Prasse for the generous provision of their lab devices, and Regina Engelhardt for excellent histopathological support.

## Funding

Supported by Else Kröner-Fresenius Foundation (EKFS, 2021_EKEA.16 and 2020_EKSP.78), CORE100Pilot Advanced Clinician Scientist Program of Hannover Medical School funded by EKFS and the Ministry of Science and Culture of Lower Saxony, and by German Center for Lung research (FKZ 82DZL002B1 & FKZ 82DZL002C1, all to J.C.S.).

## Author contributions

J.C.S. conceptualized, acquired funding, and supervised the study. J.G.: performed cohort characterization and phenotyping. L.N., F.L., D.J., F.I., J.S., J.C.K., M.K. procured, processed and characterized RAS and control specimens. L.M.L., and L.C. dissociated the lungs and isolated the nuclei. M.B. performed FANS. L.M.L., L.C., and J.C.S. performed snRNAseq bardcoding and library construction. L.M.L., L.C. and the Research Core Unit Genomics (RCUG) of MHH conducted quality control of the libraries. B.H., A.K.B. performed sequencing. All sequencing data were processed, curated, visualized, analyzed by J.C.S. and L.M.L. HE, IHC and IHF was performed by L.M.L. and analyzed by L.M.L, J.C.S and D.J. Histologic images were generated by L.M.L. L.C. prepared the samples for micro-CT experiments which were scanned by J.R., L.K., and analyzed by J.R., L.K., J.C.S., L.M.L.; E.K.J.P. created the graphical abstracts based on specifications from J.C.S. and L.M.L.; M.G., A.J., U.M., A.Ö.Y., B.V., C.F., N.K. provided critical interpretation, review, and commentary of data and manuscript. The manuscript was drafted by L.M.L. and J.C.S. and was reviewed and edited by all authors.

## Conflict of interests

J.C.S. – personal fees from Boehringer Ingelheim, MSD, Kinevant, and research grant from Deutsche Forschungsgemeinschaft (DFG), German center of Lung Research (DZL), Thyssen foundation, Else Kroener-Fresenius Foundation, Ann Theodore Foundation; J.G. - Institutional grants from DFG and DZL, consultancy fees from Zambon and Sanofi. J.G. served as a member of the CAN CLAD Study data safety monitoring board; A.J. – grants from ANR – MLQ-CT and Fondation du Souffle; F.I. – Personal fees and grants from Biotest AG and Travel fees from Biotest AG, Xvivo, Abbott. C.W. - Personal fees from Boehringer Ingelheim.

## Data and materials availability

Transcriptomic data will be deposited to the GEO, the computational code to Github.

## SUPPLEMENT

### Methods

#### Selection of RAS and Control specimens for nuclei

All CLAD specimens available at the biobank of Medical School Hanover (MHH) are phenotyped according to the latest ISHLT guidelines.(5) Formalin-fixed peripheral lung tissue from the upper lobe from 15 explant lungs from the department of pathology at MHH were used.

#### Nuclei isolation for Chromium Fixed RNA Profiling for Multiplexed Samples

Under a biological hood, each specimen was transferred in a plastic petri dish on ice and was cut with a scalpel into tiny pieces (∼1 mm³). Specimen were resuspended in 1 mL Tissue Resuspension Buffer (0.496X PBS (1X PBS, pH 7.4, Gibco) + 50 mM Tris buffer, pH 8.0 (Tris (1 M), pH 8.0, RNase-free, Invitrogen) + 0.02 % BSA (BSA, Molecular Biology Grade, NEB) + 0.24 U/µL RNase Inhibitor (Protector RNase-Inhibitor, Sigma-Aldrich) + fill up to 1 mL with Nuclease-free water (Invitrogen) on ice. Specimens were centrifuged at 850 rcf for 5 min at RT using a swinging-bucket rotor. Supernatants were removed without disturbing the tissue pellets. Meanwhile, 2 mL/specimen of Tissue Digestion Buffer (250 µg/mL Liberase TM (Liberase™ TM Research Grade, Roche) + 2.5 mg/mL Collagenase D (Collagenase D from Clostridium histolyticum, Roche) + 0.5 mM CaCl2 (Calcium Chloride Solution 2.5 M, Jena Bioscience) + fill up with 1X PBS (1X PBS, pH 7.4, Gibco™) or 10X PBS (final conc. 1X) (PBS - Phosphate-Buffered Saline (10x), Invitrogen™) + Nuclease-free H2O (Nuklease-freies Wasser (nicht DEPC-behandelt), Invitrogen™)) was pre-warmed for 10 min at 37 °C in a water bath. 2 mL pre-warmed Tissue Digestion Buffer was added to each sample. Regarding manual dissociation, a gentleMACS Octo Dissociator with heaters was used and the following program was set: Incubation for 45 min at 50 rpm, 37 °C, followed by spinning for 30 sec at (+)2000 rpm, 37°C and 30 sec at (-)2000 rpm, 37 °C. Before running the program, it was confirmed that no tissue pieces were attached to the C-tubes’ walls but all tissue pieces were resuspended in Tissue Digestion Buffer. C-tubes were detached and centrifuged at 300 rcf for 30 sec (swinging-bucket rotor) to collect all tissue pieces at the bottom of the C-tubes. Each pellet was resuspended in its supernatant using wide-bore tips. In order to remove undissociated tissue pieces and debris, each specimen was passed through a 70 µm filter (pluriStrainer® 70 µm (Zellsieb), pluriSelect) into a pre-cooled 50 mL falcon. Each filter and C-tube was rinsed with a total of 2 mL PBS. Filtrates were collected in the corresponding 50 mL falcons. Specimens were centrifuged at 850 rcf for 5 min at 4 °C using a swinging-bucket rotor. Supernatants were removed without disturbing the pellets. Each pellet was resuspended in 1 mL Quenching Buffer. To increase the relative ratio of nuclei in the suspension, specimens were filtered into 1.5 mL LoBind tubes using 20 µm filters (pluriStrainer Mini 20 µm (Zellsieb), pluriSelect). Nuclei concentration was determined using a LUNA-FL™ Dual Fluorescence Cell Counter and Acridine Orange/Propidium Iodide Stain (AO/PI) both by Logos Biosystems. Enhancer was warmed at 65 °C for 10, then briefly vortexed and centrifuged. 0.1 x specimen’s volume of pre-warmed Enhancer was added to each specimen followed by gently pipette mixing the specimens. Specimens were stored at 4 °C in the fridge for up to one week.

#### Fluorescence Activated Nuclei Sorting (FANS) of DAPI+ single nuclei

Nuclei were sorted at the research facility Cell Sorting (‘Sorter-Laboratory’) at MHH. For flow cytometric sorting of nuclei specimens were stained with 4′,6-diamidino-2-phenylindole (DAPI, Thermo Scientific™) immediately before sorting. The final concentration of DAPI was between 1 and 10 µg/mL and was adjusted if necessary to achieve saturated staining of the nuclei. The sorting of nuclei was carried out in the Central Research Facility Cell Sorting of the MHH on cell sorters equipped with a 100 µm nozzle at a pressure of 35 psi (FACSAria III Fusion, FACSAria IIu Becton-Dickinson, or MoFlo XDP, Beckman-Coulter). 5 x 105 DAPI+ single nuclei per specimen were sorted into individual 15 mL tubes , each containing 1 mL of 0.5X PBS + 0.02% BSA. Intact single nuclei were sorted by flow cytometry based on light scattering characteristics and DNA content, measured as the signal of the fluorescent DNA dye DAPI. A narrow gate in the DAPI channel was set to include only intact nuclei and to exclude debris (Fig. S1A). In further gating steps, doublets were excluded using FSC pulse parameters (Fig. S1B) and the main population was further narrowed down in a FSC-SSC gate (Fig. S1C). Reanalysis of 300 sorted nuclei from each specimen showed a homogeneous population of nuclei (Fig. S1D). Sorted single nuclei suspensions were stored on ice and were immediately used for downstream analyses as soon as 16 specimens were sorted (Chromium Fixed RNA Profiling Reagent Kits for Multiplexed Samples).

#### 10x Genomics Chromium Fixed RNA Profiling for Multiplexed Samples

Sorted DAPI+ single nuclei were immediately further processed according to “User Guide, CG000527, Rev B, Chromium Fixed RNA Profiling Reagent Kits for Multiplexed Samples” including the following choices and deviations:

Multiplexing experiment design “Maximize Number of Cells” without sub-pooling and post-hybridization pooled wash workflow was chosen for all specimens. Hybridization was performed in PCR tube-strips. Due to upfront sorting of 500,000 DAPI+ single nuclei per specimen and in order to minimize nuclei loss, the first counting step during post-hybridization was not performed. Further, all specimens (16 specimens/pool) were completely pooled because the absolute number of nuclei was estimated to be still nearly the same across all specimens due to FANS. Before passing the sample pool through a 20 µm filter (pluriStrainer Mini 20 µm (Zellsieb), pluriSelect), the nuclei pellet was only resuspended in 25 % of the volume stated in the protocol to achieve a high nuclei concentration without having to concentrate nuclei suspension after counting them (step “Cell Suspension Volume Calculator for Multiplexing 16 Samples”). 128000 nuclei were chosen as “Targeted Cell Recovery” but a 20 % higher volume of nuclei suspension stock/reaction than stated was used for the previous determined nuclei concentration and was filled up to 40 µL with Post-Hyb Resuspension Buffer. For the Sample Index PCR, a total of 12 cycles was appropriate. Three different QC-approaches per library were performed for post library construction: First a 1:20 dilution of stock library and Buffer EB (Buffer EB, Qiagen) was prepared of which 5 µL were used to determine a Qubit quantification of the 1:20 diluted library. Therefore, Qubit™ 1X dsDNA Assay-Kits (HS) + (BR) (Invitrogen™), Qubit™ assay tubes (Invitrogen™) and a Qubit™ 4 Fluorometer (Invitrogen™) were used. Afterwards, 5 µL of each 1:20 diluted library were forwarded to the Research Core Unit Genomics (RCUG) at MHH to perform a DNA fragment length analysis (HS) and a Qubit quantification. Additionally, a qPCR was performed for each library using the KAPA Library Quantification Kit for Illumina® Platforms by Roche (KR0405 – v11.20) following their detailed protocol. Remaining stock libraries and 1:20 diluted libraries were stored at -20 °C.

#### Sequencing; data processing and analysis

cDNA libraries were sequenced on an Illumina NovaSeq 6000 plattform at the Department of Human Genetics (Hanover Medical School, Germany), aiming for 800 million reads per library with a sequencing configuration of 28 base pairs on read1, 90 base pairs on read2, and 10 base pairs on Index5 and Index7, respectively. Basecalls were converted to reads using the software Cell Ranger (v7.1.0) and its bcl2fastq implementation “mkfastq”.

Read processing. Subsequent read processing was conducted using the the software Cell Ranger (v7.1.0). Reads of probes were aligned to the human reference genome GRCh38 (GENCODE v32/Ensembl 98, “GRCh38-2020-A”) using the Chromium Human Transcriptome Probe Set _v1.0.1 for GRCh38-2020-A. Technical summaries on sequencing and data processing can be accessed in supplemental table S2.

Dataset integration. Dataset integration was performed as described previously(21). Nuclei-UMI count matrices were analyzed using the R package Seurat (version 4.3.0). UMI counts were normalized with a scale factor of 10,000 UMIs per nuclei and then natural log transformed using a pseudocount of 1. To discern structural lung cells from immune cells, a recursive process of integration, graph embedding, and cluster analysis was employed. Each iteration involved integrating subjects and clustering, following Seurat’s recommended approach utilizing reciprocal PCA (https://satijalab.org/seurat/articles/integration_rpca.html). Initially, the top variable genes per subject were identified through Seurat’s FindVariableGenes using the “vst” parameter. Patterns of shared variance in these genes in respective subject were then utilized to integrate the datasets using Seurat’s FindIntegrationAnchors and IntegrateData functions, followed by scaling the resulting integrated expression matrix with Seurat’s ScaleData.

Dimension reduction, graph embedding, clustering and visualization. Dimension reduction for visualizing and clustering was performed in a stepwise process, as performed previously (13,21). Scaled values from the integration approach underwent principal component analysis (PCA), with selected principal components chosen based on their contribution to variance. Respective principal components were then used to estimate Euclidean distances between nuclei in feature-space, leading to graph embedding where edges connected the nearest neighbors for each nuclei. Louvain clustering was then applied to this network of connected nuclei. For visualization, nuclei distances and their graph embeddings underwent uniform manifold approximation and projection (UMAP). Clusters that showed no significant differences in gene expression were merged. This iterative process was continued until all nuclei from all subjects could be assigned to a discrete cell type whose characteristic features were consistently represented in all subjects. After the comprehensive categorization of all cells into different cell type clusters, we then determined the respective lineage assignment of each cluster based on the expression profile of the follwing classical lineage markers: CDH5 and PECAM1 for endothelial, MUC1 and KRT7 for epithelial, PDGFRA and PDGFRB for mesenchymal, and PTPRC for immune cells. Mesothelial were grouped together with the mesenchymal cells. As this project focused on structural cells of the RAS lungs, immune cells were excluded from downstream analyses. Multiplets and other cell barcodes of subpar quality were flagged based on their transcriptomic profiles resembling a combination of two or more distinct cell type signatures already present in the dataset. This scrutiny extended to the entire dataset and all lineage subsets. Barcodes flagged as multiplets or of low quality were excluded from subsequent analyses.

Identification of cell type specific marker genes. Cell-type specific marker genes were identified using the Wilcoxon rank sum test by comparing all cells within a specific cluster to all other cells, as implemented by Seurat’s FindMarkers function. All p-values were adjusted for multiple testing using the Bonferroni correction. An adjusted p-value below 0.05 was considered significant.

#### Preparation of formalin fixed paraffin embedded (FFPE) specimens for histologic stainings

Tissue specimen as provided by the pathology department of MHH were put into tissue cassettes. After incubating tissue for 24 h in 10% neutrally buffered formalin, specimens were stored in 70% ethanol. Then, specimens were embedded in paraffin by members of MHH’s pathology department or by LML using a tissue embedding center and stored at RT. Our standardized cutting thickness for all histologic stainings in this study is 3 µm. Cut FFPE specimens were transferred onto Superfrost™Plus Adhesion Microscope Slides, dried and stored at RT.

#### Primary antibodies

All primary antibodies were tested multiple times upfront on suitable tissue types using IHC and IHF to guarantee accurate staining. The following primary antibodies were used for our IHF (and EvG-IHC) stainings of RAS and control specimens:

| Target | Species | Clonality | Clone | Conjugate | Manufacturer | REF | Dilution IHC | Dilution IF |
| --- | --- | --- | --- | --- | --- | --- | --- | --- |
| AGER | Mouse | mono | A11 |  | Santa Cruz | sc-80652 | 1:200 | 1:25 |
| Cathepsin E (CTSE) | Rabbit | poly |  |  | Sigma Aldrich | HPA012940 | 1:200 | 1:100 |
| CD31 | mouse | mono | JC70A |  | Agilent Technologies Sales (DAKO) | M082329-2 | 1:100 | 1:50 |
| COL15A1 | Rabbit | poly |  |  | thermofisher | PA553667 | 1:600 | 1:300 |
| COX2 | Mouse | mono | COX229 |  | thermofisher | 35-8200 | 1:100 | 1:50 |
| ITGAVB6 | rabbit | mono | EM05201 |  | Sigma Aldrich | ZRB1104 | 1:100 | 1:50 |
| KRT17 | mouse | mono | E-4 | AF647 | Santa Cruz | sc-393002<br>AF647 | 1:50 | 1:25 |
| LAMP3 | goat | poly |  |  | biotechne/R&D systems | AF4087 | 1:200 | 1:200 |
| p63 | Mouse | mono | A4A |  | abcam | A8735-100 l | 1:100 | 1:50 |
| PDPN | Mouse | mono | D2-40 | AF647 | biolegend | 916606 | 1:1000 | 1:50 |
| PLVAP | Rabbit | mono | E3X9D |  | Cell Signaling Technology | 38238 | 1:100 | 1:50 |
| PRX | Rabbit | poly |  |  | novus bio | NBP1-89598-25ul | 1:400 | 1:200 |
| SFTPC | Mouse | mono | H8 |  | Santa Cruz | sc-518029 | 1:200 | 1:100 |

#### Secondary antibodies for IHF stainings

All secondary antibodies were tested multiple times upfront in combination with reliable primary antibodies. The following antibodies were used for our IHF stainings:

| Target | Species | Conjugate | Manufacturer | REF | Dilution |
| --- | --- | --- | --- | --- | --- |
| Goat-IgG | Donkey | AF488 | Biozol | JIM-705-545-147 | 1:200 |
| Mouse-IgG | Horse | DyLight 488 | VectorLabs | DK-8828 | ready to use |
| Mouse-IgG | Donkey | AF488 | Biozol | JIM-715-545-150 | 1:200 |
| Rabbit-IgG | Horse | DyLight 594 | VectorLabs | DK-8828 | ready to use |
| Rabbit-IgG | Donkey | Alexa 647 | Jackson ImmunoResearch | 711-607-003 | 1:200 |
| Rabbit-IgG | Donkey | Rhodamine Red™-X (RRX) | Jackson ImmunoResearch | 711-295-152 | 1:200 |

#### Immunohistochemistry (IHC)

IHC was performed to test the functionality and accuracy of primary antibodies. Before starting, all reagents were equilibrated to RT. A standard deparaffinization and rehydration of FFPE tissue was performed (2x 5 min xylene, 2x 3 min 100% ethanol, 1x 3min 95% ethanol, 1x 1 min 70% ethanol, 1x 1 min 50% ethanol, 1x 5 min distilled water), followed by heat-induced antigen retrieval. Therefore, tissue slides were boiled for 20 min at 98 °C in 1× tris-based Antigen Unmasking Solution (Vector Laboratories, USA), diluted 1:100 with distilled water. After letting specimens cool for 20 min, they were washed in 1x PBS for 5 min. Tissue sections were encircled with a Roth®-Liquid Barrier Marker and put into a wet chamber which was used to store slides during all further incubation steps. Specimens were incubated for 10 min in BLOXALL Endogenous Peroxidase and Alkaline Phosphatase Blocking Solution (Vector Laboratories, USA) in order to block endogenous peroxidase and alkaline phosphatase activity. Tissue slides were washed for 5 min in 1x PBS and blocked for 20 min using 2.5% Normal Horse Serum Blocking Solution (Vector Laboratories, USA). Each specimen was incubated for 30 min with a primary antibody (diluted appropriately in 2.5% Normal Horse Serum Blocking Solution) at RT. After washing the tissue slides in 1x PBS for 5 min, each specimen was incubated for 30 min with a corresponding secondary antibody (anti-mouse, anti-rabbit or anti-goat ImmPRESS reagent, conjugated with horseradish, all Vector Laboratories, USA) at RT. Slides were washed once again for 5 min in 1x PBS, then incubated for 6-10 min in ImmPACT® DAB Substrate Kit, Peroxidase (HRP) (Vector Laboratories, USA). Incubation time varied due to each individual reaction times. After one last wash for 5 min in 1x PBS, specimens were counterstained in Hematoxilin Solution Gill no. 1 (Sigma-Aldrich, USA) for 3, then washed for 15 min in tap water (intermittent exchange of tap water). Afterwards, a standard dehydration in ethanol/xylene was performed (1x 15 sec 50% ethanol, 1x 15 sec 70% ethanol, 1x 1 min 95% ethanol, 2x 1 min 100% ethanol, 2x 3 min xylene). Tissue slides were mounted with VectaMount permanent mounting solution (Vector Laboratories, USA) and coverslips were put on. Stained slides were digitalized and analyzed using a ZEISS Axioscan 7 and the ZEN 3.5 (blue edition) software. Slides were stored at RT.

#### Immunhistofluorescent (IHF) stainings

Before starting, all reagents were equilibrated to RT. A standard deparaffinization and rehydration of FFPE tissue was performed (2x 5 min xylene, 2x 3 min 100% ethanol, 1x 3min 95% ethanol, 1x 1 min 70% ethanol, 1x 1 min 50% ethanol, 1x 5 min distilled water), followed by heat-induced antigen retrieval. Therefore, FFPE tissue slides were boiled for 20 min at 98 °C in 1× tris-based Antigen Unmasking Solution (Vector Laboratories, USA), diluted 1:100 with distilled water. Afterwards specimens were cooled down for 20 min in 1X PBS at RT. Meanwhile, a stock of 0.9995X PBS + 0.05% Tween® 20 was prepared and used to prepare 2.5% normal serum (donkey/goat/horse) corresponding to the secondary antibodies’ species. All primary and secondary antibodies were diluted in suitable 2.5% normal serum and stored at RT under tin foil until needed. After letting the slides cool down, they were encircled with a Roth®-Liquid Barrier Marker and put into a wet chamber which was used to store slides during all further incubation steps. Specimens were incubated for 20 min with 2.5% normal serum before gently tapping the slides sideways on paper towels to remove normal serum. No washing step was performed at this point. Each specimen was incubated for 60 min with the prepared solution of 2.5% normal serum and primary antibody/antibodies at RT. Slides were washed for 2x 3 min in PBS containing 0.05% Tween® 20. Specimens were incubated for 60 min with the prepared solution of 2.5% normal serum and secondary antibody/antibodies under tin foil at RT. Slides were washed in staining throughs covered with tin foil for 2x 3 min in PBS containing 0.05% Tween® 20. Specimens were incubated for 60 min with the prepared solution of 2.5% normal serum and fluorochrome conjugated antibody/antibodies under tin foil at RT. Slides were washed in staining throughs covered with tin foil for 2x 3 min in PBS containing 0.05% Tween® 20. Vector TrueView Autofluorescence Quenching Kit (Biozol, VEC-SP-8400-15) was used according to the manufacturer’s information before washing slides in staining throughs covered with tin foil for 5 min in PBS containing 0.05% Tween® 20. Slides were mounted with VECTASHIELD Vibrance™ with DAPI Antifade Mounting Medium (VectorLaboratories/ Biozol, VEC-H-1800) and coverslips were put on. Stained slides were digitalized and analyzed using a ZEISS Axioscan 7 and the ZEN 3.5 (blue edition) software. Slides were stored darkly at 4 °C.

#### Hematoxylin-eosin (HE) stainings

Before starting, all reagents were equilibrated to RT. A standard deparaffinization and rehydration of FFPE tissue was performed (2x 5 min xylene, 2x 3 min 100 % ethanol, 1x 3min 95 % ethanol, 1x 1 min 70 % ethanol, 1x 1 min 50 % ethanol, 1x 5 min distilled water). Slides were incubated with 2:1 Hematoxylin solution Gill no. 1 (Sigma, GHS-1-16) for 5 min. Afterwards, slides were washed for 15 min in tap water (intermittent exchange of tap water)., then rinsed for 2 min with distilled water. Slides were incubated for 3 min in a ready-to-use solution (Roth, X883.1), then shortly rinse with distilled water. Afterwards, a standard dehydration in ethanol/xylene was performed (1x 15 sec 50% ethanol, 1x 15 sec 70% ethanol, 1x 1 min 95% ethanol, 2x 1 min 100% ethanol, 2x 3 min xylene). Tissue slides were mounted with VectaMount permanent mounting solution (Vector Laboratories, USA) and coverslips were put on. Stained slides were digitalized and analyzed using a ZEISS Axioscan 7 and the ZEN 3.5 (blue edition) software. Slides were stored at RT.

#### RNAscope

RNAscope was performed strictly according to the “RNAscope Multiplex Fluorescent Reagent Kit v2 User Manual” (UM 323100/Rev B/ Effective Date: 10/11/2022) by ACD bio. Slides were stored at 4°C in the dark until scanning. Stained slides were digitalized and analyzed using a ZEISS Axioscan 7 and the ZEN 3.5 (blue edition) software.

List of chemicals:

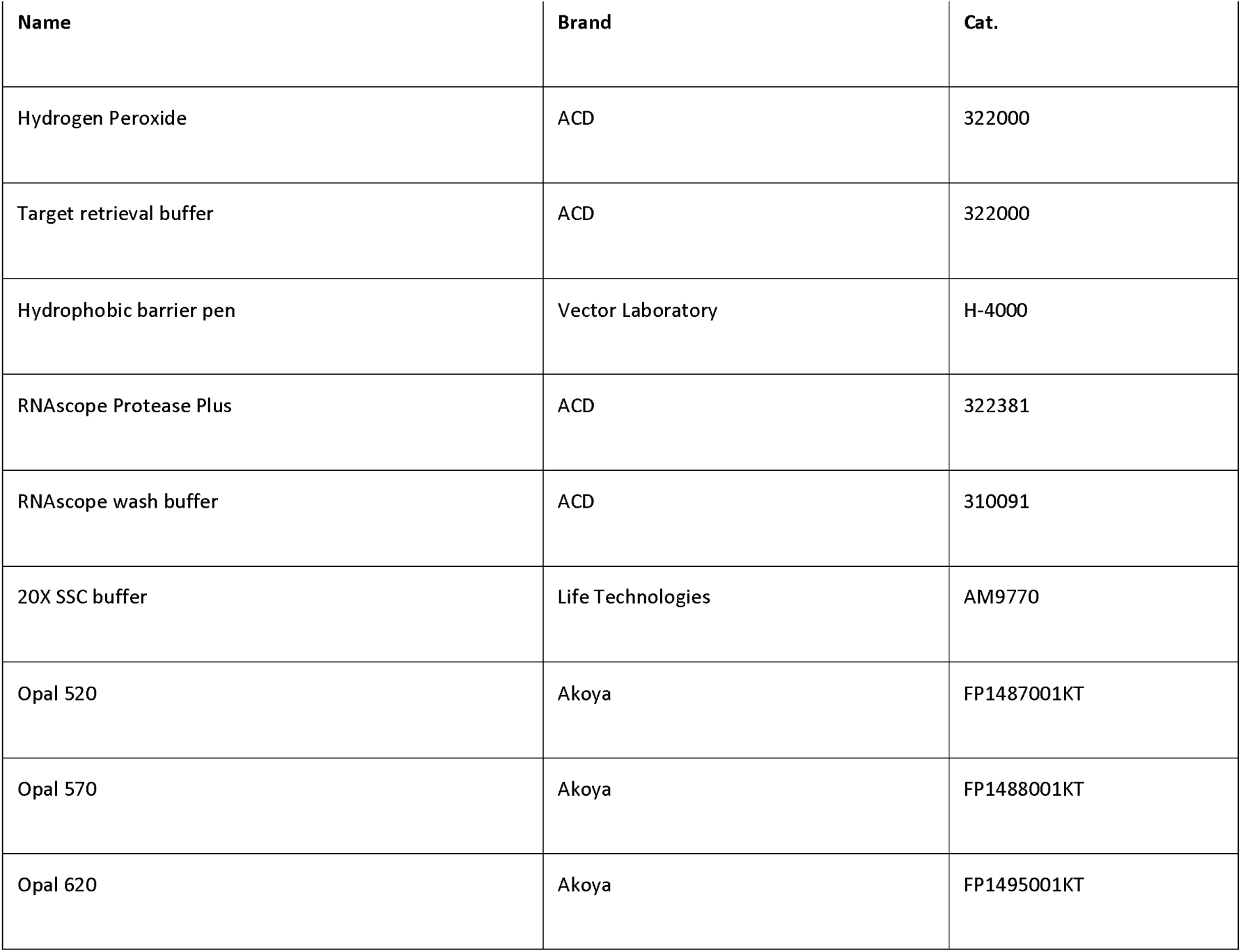
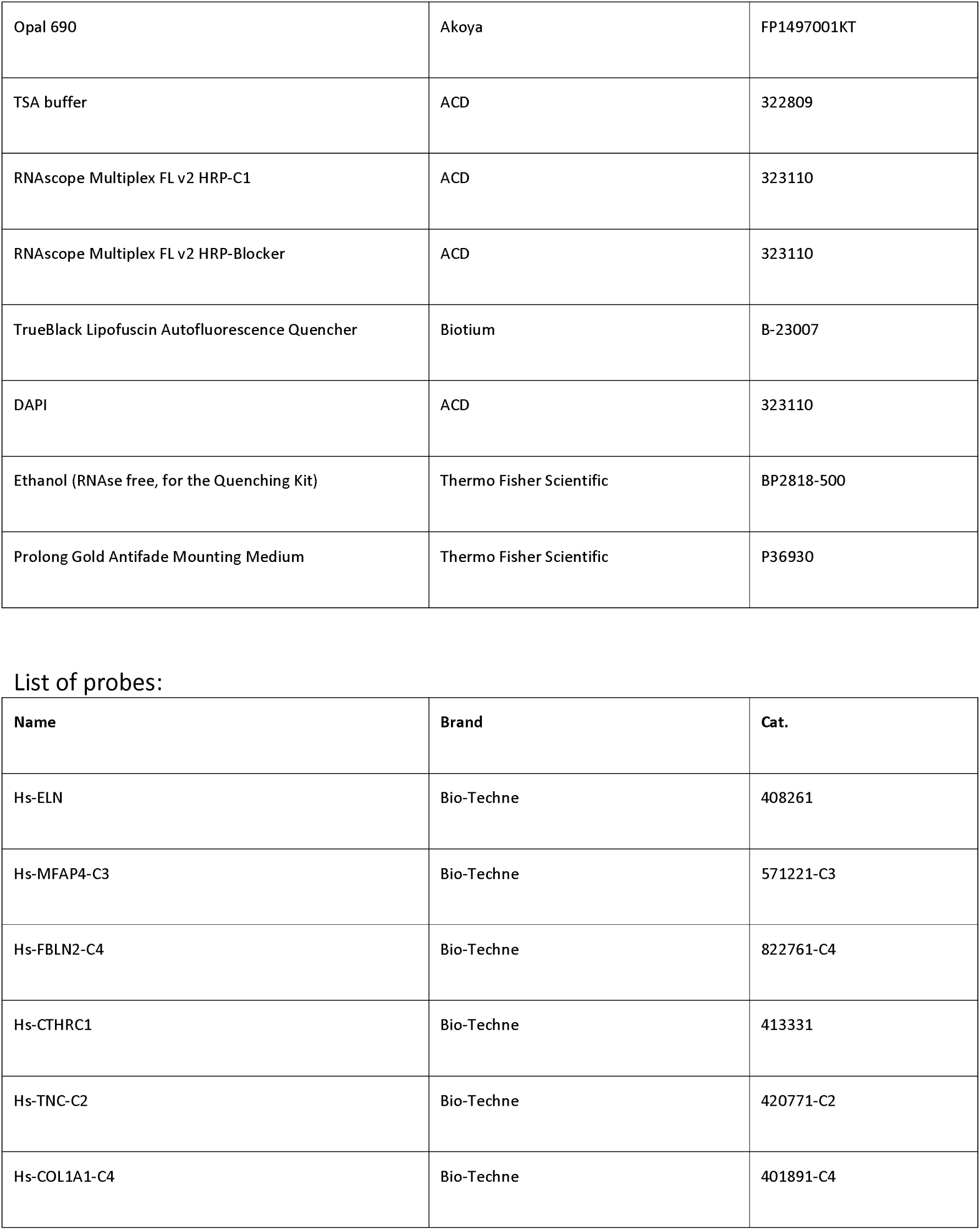

#### Elastica van Gieson (EvG) stainings

A standard deparaffinization and rehydration of FFPE tissue was performed (1:30 min xylene, 1:30 min 100% ethanol, 1:30 min 90% ethanol, 1:30 min 70% ethanol). Afterwards, specimens were put for 10 min in Resorcinol-Fuchsine-solution acc. to Weigert, then differentiated for 2x 0:30 min in 100% ethanol. Specimens were for 10 min with iron hematoxylin solution acc. to Weigert, then blued for 5 min in tap water. Slides were stained for 0:15 min in Van Gieson’s solution (picrofuchsin) before dehydrating them (0:05 min 70% ethanol, 1:30 min 90% ethanol, 2x 1:30 min 100% ethanol, 1:30 min xylene). Stained slides were digitalized and analyzed using a ZEISS Axioscan 7 and the ZEN 3.5 (blue edition) software. Slides were stored at RT.

#### Combined Elastica van Gieson (EvG) and Immunohistochemistry (IHC) stainings

A standard deparaffinization and rehydration of FFPE tissue was performed (2x 10 min xylene, 2 min 100% ethanol, 2 min 90% ethanol, 2 min 70% ethanol, 2 min 50% ethanol, 2 min demineralized water). Then, tissue slides were boiled for 30 min at 98°C in Tris pH 9.0, and washed for 10 min in demineralized water at RT. Tissue sections were encircled with a Roth®-Liquid Barrier Marker. Antibody staining was performed as following: 5 min 3% H_2_O_2_, 5 min washing buffer, 5 min blocking solution, 5 min washing buffer, 1h antibody incubation, 5 min washing buffer, 30 min HRP-polymer, 5 min washing buffer, 15 min DAB (1:20 diluted), and 5 min washing in demineralized water. Slides were air-dried. Afterwards, EvG stainings were automatically performed according to the EvG staining protocol stated above (but without the deparaffinization/rehydration), using a HistoCore SPECTRA Workstation (Workstation-Kit - 14051254355, ST - 14051254354, CV – 14051454200).

#### Micro-CT

RAS specimen were contrasted using Tungsten phosphoric acid for 48h, then scanned with a Phoenix Nanotom® M micro-CT scanner (Waygate technologies). A voxel size of 8.46 µm was acquired employing a x-ray tube potential (peak voltage) of 60kV given a x-ray intensity current of 540 µA. Micrographs were obtained with an average of 5489 distinct grey scales securing optimized optical contrast. Raw data image processing applying a ROI- and Inline Median was performed prior to final 3D volume rendering, which was conducted with VGSTUDIO 2022 64-Bit Software (Volume Graphics).

#### Image processing

All histologic images were analyzed and processed using ZEN (blue edition) software by ZEISS. The processing method “background subtraction” was applied to all IHF and RNAscope images using default settings. White and black values were manually adjusted for each channel. No further image processing methods were used. Multi-panel figures were put together using Adobe Illustrator 2024.

## SUPPLEMENTAL TABLE and TABLE LEGENDS

**Supp. Table 1:** Patient characteristics. Basic characteristics of all included controls and RAS patient. Continuous variables are depicted as median and interquartile range. Age is given in years. FVC, FEV1 and TLC are calculated as percent of the baseline values, the mean of the best two post-operative measurements. All lung function values show last available lung function pre re-transplantation, FEV1=Forced expiratory volume in 1 second; FVC= Forced vital capacity; n.a.=not applicable; TLC=total lung volume.

| n | Control<br>n=9 | RAS<br>n = 15 |
| --- | --- | --- |
| Female / Male [n] | 4/5 | 6/9 |
| Downsizing / Tumor-distant tissue [n] | 5/4 | n.a. |
| Age in years | 57 [54-63] | 44 [35-55] |
| FEV1 % baseline | n.a. | 27 [17-37] |
| FVC % baseline | n.a. | 41 [38 - 46] |
| TLC % baseline | n.a. | 74 [71 - 86] |
| Immunosuppressive Regime | n.a. | Tacrolimus (n=11),<br>Ciclosporine (n=3),<br>Prednisolone/mycophenolate n=15 |

**Supp. Table 2:** see separate Excel table

**Legend Supp. Table 2: 2A)** Summary of technical summaries of read processing for per sample. **2B**) Full cohort marker table. Results of Wilcoxon rank-sum test of each cell-type in the whole dataset against the other varieties in the whole dataset per cell type. **2C**) Epithelial subset marker table. Results of Wilcoxon rank-sum test of each epithelial cell-type against the other epithelial varieties per cell type. **2D**) Endothelial subset marker table. Results of Wilcoxon rank-sum test of each endothelial cell-type against the other endothelial varieties per cell type. **2E**) Stromal subset marker table. Results of Wilcoxon rank-sum test of each stromal cell-type against the other stromal varieties per cell type. **2F)** Results of Wilcoxon rank-sum test comparing pathway scores of each cell-type in the whole dataset against the other varieties in the whole dataset per cell type. **2G)** RAS versus control differential expression table. Results of Wilcoxon rank-sum test per cell type comparing RAS to control. **2H)** Cell differential statistics. Frequencies and results of the Wilcoxon rank-sum test comparing of RAS versus control cell-type makeup as a fraction of their respective lineage group.

## SUPPLEMENTAL FIGURE LEGENDS

**Figure S1:** (A) A narrow gate in the DAPI channel was set to include only intact nuclei and to exclude debris. (B) In further gating steps, doublets were excluded using FSC pulse parameters. (C) The main population was further narrowed down in a FSC-SSC gate. (D,E) Reanalysis of the sorted nuclei showed a homogeneous population of nuclei, which were used for downstream analyses.

**Figure S2: A:** IHF stains of control specimens. Scale bars are 100 µm. Arrows A (insert arrow) indicate *KRT17+*/*TP63+* basal cells. **B**: EvG stains of control specimens. Scale bars are 100 µm.

**Figure S3:** IHF stains of control specimens. Scale bars are 100 µm. Arrows A (insert arrow) indicate *COL15A1+/PLVAP+* peribronchial VE cells. Arrow B (insert arrow) indicates *PRX+* microvasculature. **B**: IHC-EvG stainings of control specimens. Scale bars are 100 µm. Arrows indicate *CD31+* VE cells and *COL15A1+* peribronchial VE cells.

